# Engineering BCMA CAR T cells for myeloma-targeted cargo delivery

**DOI:** 10.1101/2024.02.14.580301

**Authors:** Thomas Kimman, Marta Cuenca, Anne Slomp, Ralph G Tieland, Dedeke Rockx-Brouwer, Sabine Heijhuurs, Angelo D. Meringa, Wendy Boschloo, Douwe MT Bosma, Sanne Kroos, Vania lo Presti, Stefan Nierkens, Niels Bovenschen, Jürgen Kuball, Monique C Minnema, Zsolt Sebestyén, Victor Peperzak

## Abstract

Clinical responses with chimeric antigen receptor (CAR) T cells are encouraging, however, primary resistance and relapse after therapy prevent durable remission in a large fraction of cancer patients. One of the underlying causes comprises apoptosis resistance mechanisms in cancer cells that limit killing by CAR T cells. Therefore, we developed a technology that boosts tumor cell apoptosis induced by CAR T cells. We reveal that B cell maturation antigen (BCMA) CAR T cells equipped with a granzyme B-NOXA fusion construct improves killing of multiple myeloma (MM) cells, both in vitro and in a xenograft mouse model, by localizing NOXA to cytotoxic granules that are released into cancer cells upon contact. Since MM cells critically depend on MCL-1 expression, inhibition by its natural ligand NOXA effectively induces apoptosis. Overall, this strategy allows specific delivery of cargo into cancer cells and improves killing efficacy of CAR T cells in a tailor-made manner.

## Introduction

Recent clinical success of immunotherapy has made a colossal impact on the development of novel therapeutic approaches targeting cancer. Engineered T cell therapy is, together with the discovery of checkpoint inhibitors, one of the most promising types of anti-cancer immunotherapy where scientific efforts are accompanied with high investments by industry^1^. Despite their initial successes, clinical trials with engineered T cells, and specifically with chimeric antigen receptor (CAR) T cells, revealed two major weaknesses: (1) primary resistance upon treatment with CD19-specific CAR T cells in chronic lymphocytic leukemia (CLL) and diffuse large B cell lymphoma (DLBCL), and (2) disease relapse after treatment with CD19-specific CAR T cells for B cell acute lymphoblastic leukemia (B-ALL) or with B cell maturation antigen (BCMA)-specific CAR T cells for multiple myeloma (MM)^2,3^. Resolving these weaknesses could greatly increase effectivity and broad applicability of engineered T cell therapy against cancer. Previously described limitations to successful CAR T cell therapy include antigen loss on cancer cells and failed CAR T cell expansion and persistence^4^. While multiple strategies have been considered to improve gene engineered T cells, attempts to directly improve their killing capacity remain neglected^5^. If initial CAR T cell-directed killing of cancer cells can be improved, subsequent selection for antigen-negative cancer cells and CAR T cell persistence become less relevant. There is ample evidence that CAR T cell-induced apoptosis is often suboptimal and that apoptosis resistance mechanisms limit effective responses to CAR T cell therapy in various B cell malignancies. For example, increased expression of pro-survival protein BCL-2 has been observed in B lymphoma cells that survive treatment with CD19 CAR T cells and therefore entails a resistance mechanism that disables the natural killing machinery of CAR T cells^6^. In line with these findings it was shown using genome-wide CRISPR/Cas9 screening that loss of pro-apoptotic BCL-2 family protein NOXA in B lymphoma cells regulates resistance to CAR T cell therapy by impairing apoptosis of tumor cells^7^. Execution of target cell apoptosis can be mediated by release of granzymes by CAR T cells or by engaging death receptors on targeted cancer cells. Interestingly, resistance mechanisms for both pathways have been described. Two separate studies used genome wide CRISPR/Cas9 knock-out screens in B-ALL cells to reveal that death receptor TRAIL-R2 (TNFRSF10B), and downstream signaling molecules FADD, BID and CASP8, mediate sensitivity to CD19 CAR T cell killing^8,9^. In addition, we have recently shown that expression of granzyme B-inhibitor serpin B9 in DLBCL and CLL cells inhibits killing by CD19 or CD20 CAR T cells, thereby revealing another apoptosis resistance mechanism^10^. Rather than bypassing specific apoptosis resistance mechanisms, such as those described above, we explored the possibility to directly enhance the killing potential of CAR T cells to dampen primary resistance and reduce the chance of disease relapse.

## Results

BCL-2 family member MCL-1 is a key pro-survival factor for MM cells and its elevated expression is associated with chemoresistance and shorter event-free survival in MM^11,12,13^. Therefore, we examined if MCL-1 expression was also enriched in MM cells that resist killing by BCMA CAR T cells (**Fig. 1a**). Human MM cell lines NCI-H929 and L363 were co-cultured with BCMA CAR T cells and intracellular MCL-1 protein expression was subsequently measured in viable MM cells. We found that MCL-1 expression was increased in surviving MM cells after co-culture with BCMA CAR T cells compared to expression in MM cells cultured alone (**Fig. 1b,c**). Since MCL-1 can be targeted with specific small molecule inhibitors we added MCL-1-inhibitor S63845 during co-culture of MM cells with BCMA CAR T cells^14,15^. This resulted in improved killing of MM cells compared to co-culture with BCMA CAR T cells alone (**Fig. 1d,e**), including for L363 MM cells that are relatively resistant to BCMA CAR T cells. These experiments combined illustrate that BCMA CAR T cell-mediated MM cell killing can be enhanced by simultaneous targeting of MCL-1 and suggest that co-treatment may be clinically advantageous. However, we and others have shown that MCL-1 expression sustains survival of many healthy cells and tissues and co-treatment with a systemic MCL-1 inhibitor would generate undesired side-effects, precluding its use as a safe anti-cancer drug^16,17,18,19,20^. For example, it was shown that genetic deletion of *Mcl1* in mice causes lethal cardiac failure and that MCL-1 inhibition promotes apoptosis of human iPSC-derived cardiomyocytes^21,22^. Regardless of these findings, multiple pharmaceutical companies generated MCL-1-specific inhibitors for treatment of MM, acute myeloid leukemia (AML) and B cell lymphoma that were tested in phase I clinical trials. Currently, most of these clinical trials have been put on FDA-instructed or voluntary hold due to dose-related cardiac toxicity^23^. Thus, although MCL-1 is a bona fide MM target, it should be inhibited in MM cells specifically to allow safe and effective treatment. To target MCL-1 specifically in cancer cells and avoid systemic toxicity we explored the possibility of MCL-1-inhibitor delivery via CAR T cells.

**Figure 1.**
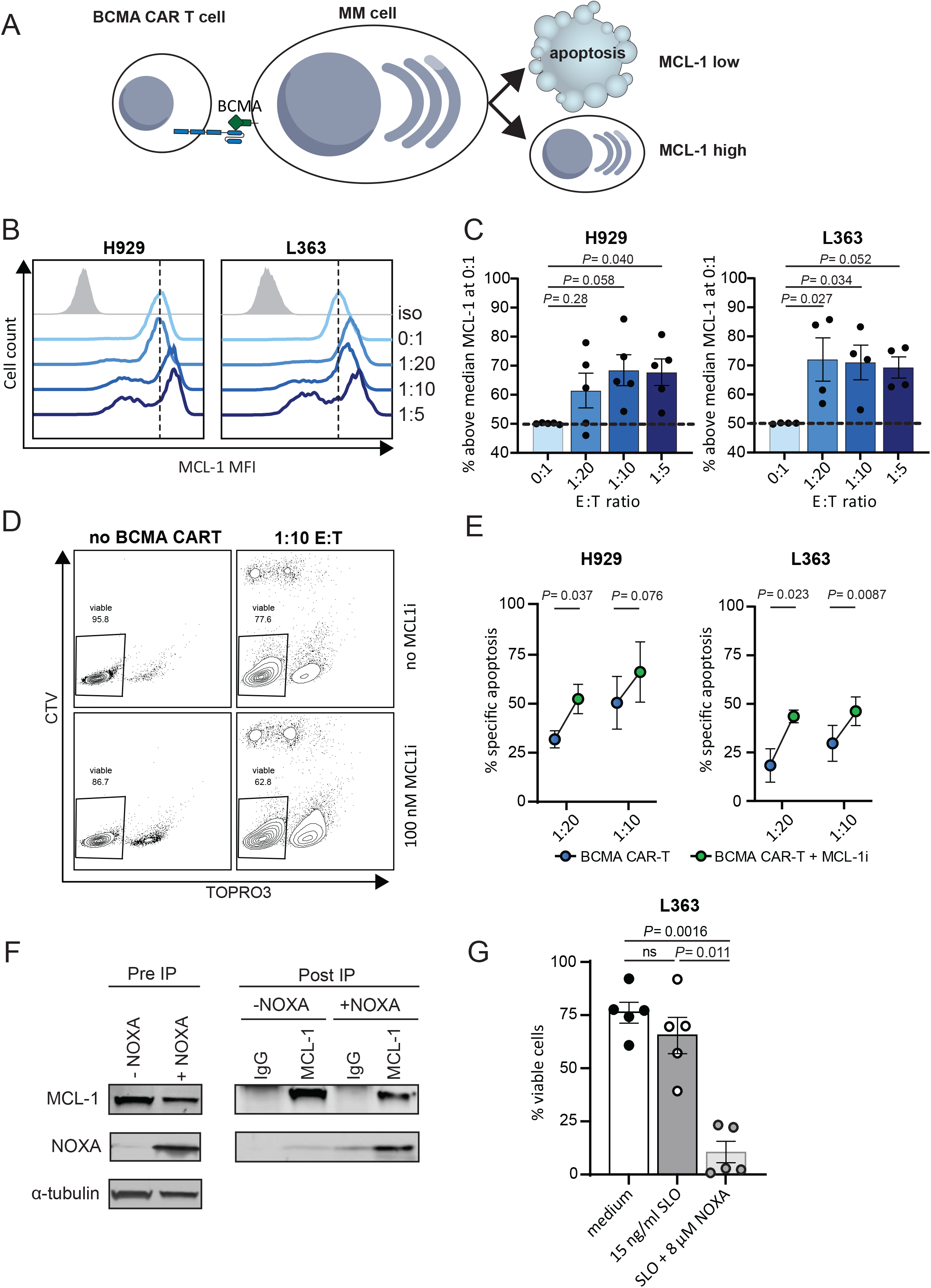
MCL-1 expression limits MM cell killing by BCMA CAR T cells. **A)** Graphical representation of experimental findings. **B**) Representative histograms showing MCL-1 protein expression in viable L363 or H929 MM cells after 24 h co-culture with BCMA CAR T cells using indicated effector to target cell ratio’s (E:T) and measured by intracellular flow cytometry. The dotted line represents the median MCL-1 expression in untreated L363 or H929 cells. Grey indicates isotype control staining. **C**) Percentage of viable H929 (n = 5) or L363 (n = 4) cells with MCL-1 expression above the median expression in untreated cells, as shown in (B). Dots represent separate experiments with SEM. Statistical testing was performed using one-way ANOVA, followed by multiple comparison testing. **D**) Representative gating strategy of L363 MM cells co-cultured with CellTrace Violet (CTV)-labelled BCMA CAR-T cells and stained with nucleic acid dye TO-PRO-3, and measured by flow cytometry after 24 h of culture. Co-cultures were simultaneously incubated with 100 nM MCL-1 inhibitor S63845 (lower panels) or without (upper panels). Indicated percentages of viable cells are calculated within CTV-negative MM cells. **E**) Quantified specific apoptosis of L363 or H929 cells as detailed for (D) and by using the gating strategy shown in (D). Percentages were calculated based on absolute cell numbers using counting beads. Specific apoptosis was determined by measuring the altered percentage of TO-PRO-3^-^ (live) cells compared with untreated cells and was defined as follows: ([% cell death in treated cells - % cell death in control]/% viable cells control) x 100. For H929 a concentration of 10 nM MCL-1i and for L363 a concentration of 100nM MCL-1i was used. Dots show averages of separate experiments with L363 (n = 6) or H929 (n = 3) with SEM. Statistical testing was performed using a paired t-test. **F**) SDS-PAGE electrophoresis of NP40 lysates from L363 MM cells treated with 15 ng/ml SLO, with or without 10 µM synthetic NOXA, stained for NOXA, MCL-1 and α-tubulin as control. Left panel shows the untreated lysates before immunoprecipitation (pre IP) and the right panel shows the cell lysates after immunoprecipitation with MCL-1 or control (IgG) antibodies (post-IP). **G**) Percentage of viable (DiOC6(3)^+^TO-PRO-3^-^) L363 MM cells treated with 15 ng/ml SLO, with or without 8 µM synthetic NOXA and analyzed by flow cytometry. Shown are averages of 5 biological replicates with SEM. Statistical analysis was performed using a one-way ANOVA with Geisser-Greenhouse correction.

BH3-only protein NOXA is a p53-inducible selective inhibitor of MCL-1 and associated with apoptosis induction in many forms of cancer, including MM^24,25^. Interestingly, it was reported that low NOXA expression in relapsed/refractory B-cell lymphoma cells correlated with worse patient survival after tandem CD19/20 CAR T cell treatment and that pharmacologically-induced expression of NOXA sensitized cancer cells to CAR T cell-mediated killing^7^. These findings indicate that the level of NOXA expression in cancer cells may determine their sensitivity to CAR T cell-mediated killing. To test whether exogenous delivery of NOXA induces apoptosis in cancer cells that depend on MCL-1 expression for survival, we treated MM cells with synthetic NOXA together with sub-lytic concentrations of pore-forming protein streptolysin O (SLO) to facilitate NOXA delivery in target cells. We could visualize entry of fluorescently labelled synthetic NOXA (NOXA-TAMRA) into the cytosol of L363 MM or OCI-Ly10 DLBCL cells by confocal microscopy (**Extended Data** Fig. 1a,b) and revealed binding to intracellular MCL-1 by immunoprecipitation (**Fig. 1f**). The introduced NOXA promoted MM cell apoptosis in a dose-dependent manner, which shows that delivery of NOXA specifically into tumor cells is sufficient to induce tumor cell killing (**Fig. 1g** **and Extended Data** Fig. 1c-e).

Next, we developed a strategy to load proteins of choice in cytotoxic granules of CAR T cells that can be released into target cells upon contact (**Fig. 2a**). The ultimate aim with this strategy is to boost CAR-T cell cytotoxicity and kill additional cancer cells that would resist killing by standard CAR T cell mechanisms, including release of cytotoxic granules containing perforin and granzymes, and ligation of death receptors on cancer cells. To achieve this, we cloned cargo proteins, including the red fluorescent protein mScarlet, behind the sequence encoding granzyme B in a lentiviral vector (**Fig. 2b**). Using confocal microscopy we confirmed that these cargo proteins localize to LAMP-1-positive cytotoxic granules in transduced primary human T cells (**Fig. 2c**). Next, we transduced BCMA CAR T cells with the construct as shown in Fig. 2b and co-cultured these with MM cells. In time, accumulation of fluorescent mScarlet could be detected in the targeted MM cells (**Fig. 2d,e**). EGFP, that was placed behind a T2A sequence and not directly located behind the granzyme B sequence, was not localized to cytotoxic granules in transduced BCMA CAR T cells and was therefore not delivered to MM cells after co-culture (**Fig. 2c,d,f**). Replacing mScarlet by the more stable fluorescent protein mNeonGreen allowed visualizing cargo transfer in timelapse confocal imaging (**Extended Data** Fig. 2a). Co-culture of BCMA CAR T cells transduced with the Granzyme B-mNeonGreen construct together with MM H929 cells showed re-localization of mNeonGreen to the synapse with MM cells, followed by delivery of a portion of the mNeonGreen cargo from the CAR T cells into the cytosol of MM cells (**Extended Data** Fig. 2b). Combined, these experiments reveal that using our strategy proteins of choice can be localized to cytotoxic granules in CAR T cells and delivered specifically into targeted cancer cells upon contact.

**Figure 2.**
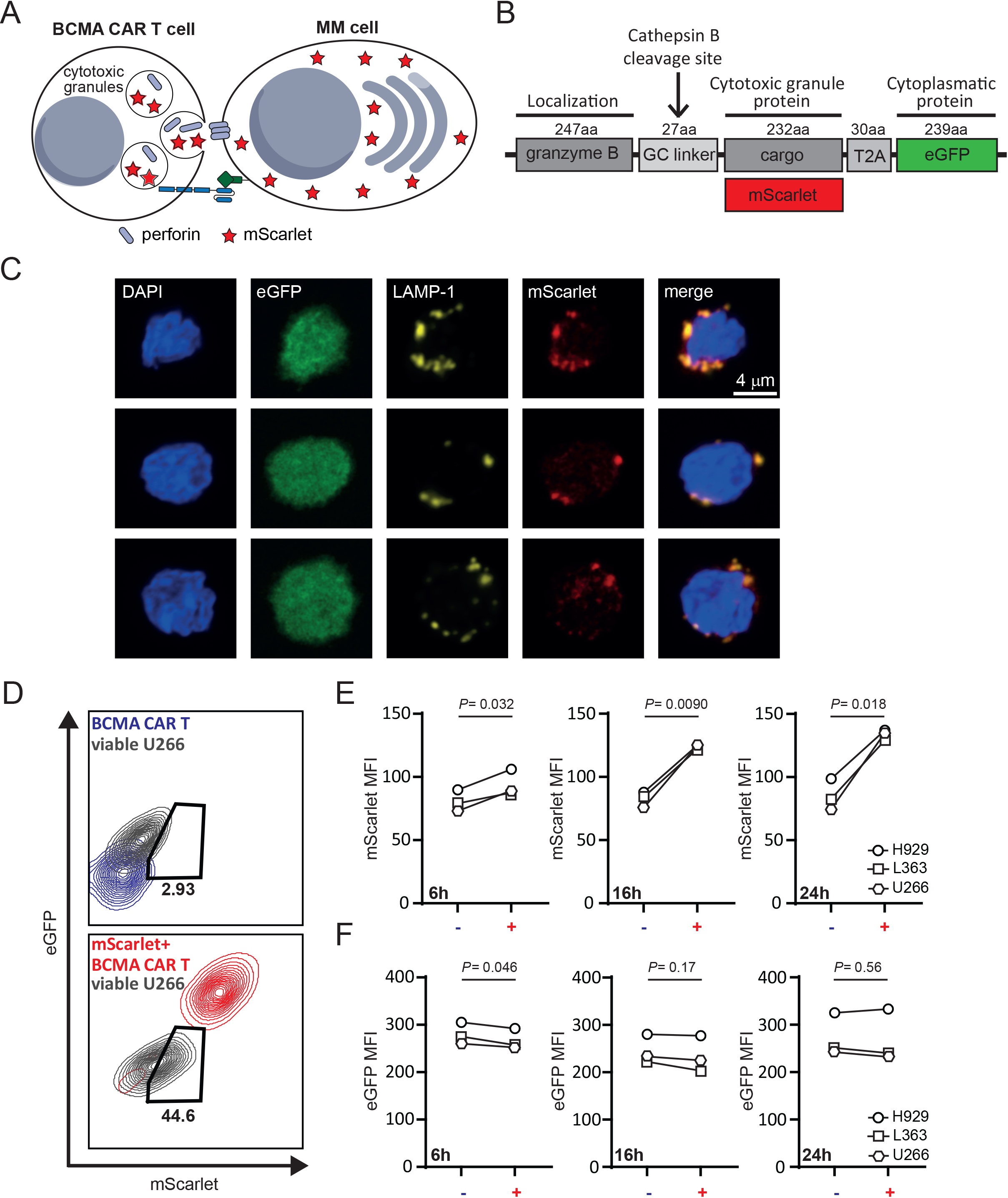
BCMA CAR T cells can be equipped with fluorescent cargo that is transferred to MM cells upon cell-cell interaction. **A)** Graphical representation of experimental findings. **B**) Construct design with cargo proteins fused to granzyme B through a GC linker that contains a cathepsin B cleavage site. **C**) Representative confocal images (630 x oil-magnification) of BCMA CAR T cells transduced with the lentiviral construct shown in (B) and stained for late endosomal marker LAMP-1 and DAPI. (**D**) Representative image of eGFP and mScarlet fluorescence in viable U266 MM cells (grey) when co-cultured with WT (blue) or mScarlet^+^ (red) BCMA CAR T cells for 24 h in a 1:5 E:T cell ratio. Fluorescence signals of WT or mScarlet^+^ BCMA CAR T cells were used as overlay in these plots to indicate range of eGFP and mScarlet expression. **E, F**) Mean fluorescence intensity (MFI) of mScarlet (E) or eGFP (F) in the total viable population of indicated target MM cell lines when co-cultured for 6, 16 or 24 h with WT (-) or mScarlet^+^ (+) BCMA CAR T cells in a 1:5 E:T cell ratio. The experiment was performed in the presence of 10 µM caspase inhibitor Q-VD-OPh to inhibit apoptosis of target cells. Statistical analysis was performed using a paired T-test.

To test improved killing of MCL-1-dependent cancer cells by CAR T cells, we cloned the sequence encoding NOXA behind granzyme B, as we did for mScarlet (**Fig. 3a,b**). As control, we introduced 3 point mutations in the BCL-2 homology 3 (BH3) domain of NOXA, which is used for binding and inhibiting MCL-1 (**Fig. 3b**). These point mutations render NOXA inactive (iNOXA) and unable to bind MCL-1^26^. By using HA-tagged versions of NOXA and iNOXA we could visualize their intracellular localization in transduced T cells. As expected from our findings with fluorescent cargo molecules (**Fig. 2c**), we observed that NOXA and iNOXA localize to LAMP-1-positive cytotoxic granules in transduced primary T cells or in NK cell line YT-Indy (**Extended Data** Fig. 3a-c). BCMA CAR T cells transduced with constructs containing NOXA (NOXA-BCMA CAR T cells) increased apoptosis of MM cell lines H929 (**Fig. 3c**) and RPMI-8226 (**Extended Data** Fig. 4a), as well as primary MM cells (**Fig. 3d**), compared to control BCMA CAR T cells transduced with iNOXA (iNOXA-BCMA CAR T cells). Our findings were not only restricted to BCMA CAR T cells since YT-Indy NK cells transduced with constructs containing NOXA killed H929 MM cells (**Extended Data** Fig. 4b) and diffuse large B cell lymphoma (DLBCL) cell line OCI-Ly7 (**Extended Data** Fig. 4c) better than NK cells transduced with iNOXA. To confirm improved killing by NOXA-BCMA CAR T cells in an in vivo model, we performed mouse xenograft experiments. Here, immune deficient NOD SCID gamma (NSG) mice were intravenously injected with luciferase-transduced RPMI-8226 MM cells. Three weeks after tumor engraftment, when tumor cells could be visualized, NOXA-BCMA or iNOXA-BCMA CAR T cells were intravenously injected (0.8 x 10^6^ CAR T cells per mouse in a 1:1 CD4:CD8 T cell ratio) and tumor growth was monitored in time using bioluminescence imaging (BLI) (**Fig. 3e**). In this in vivo model, MM outgrowth was significantly delayed in mice receiving NOXA-BCMA CAR T cells as compared to mice receiving iNOXA-BCMA CAR T cells, similar to in vitro observations (**Fig. 3f,g**). Strategies to indirectly improve CAR T cell killing efficacy have previously been demonstrated. This includes CAR T cells secreting proinflammatory cytokines IL-12 or IL-18 that influence the tumor microenvironment and potentiate the antitumor response^27,28^. With these approaches secretion is not directed specifically towards cancer cells, which may result in toxicity to healthy cells and tissues. In contrast, our optimized killing strategy for CAR T cells delivers a pro-apoptotic molecule specifically into cancer cells that interacted with a CAR T cell. Due to the directed secretion that is limited to the immune synapse, toxicity to neighboring cells is expected to be minimal. To examine this in more detail, we tested NOXA-BCMA CAR T cell-mediated toxicity to non-targeted BM cells from MM patients in co-culture experiments. Although a clear reduction in MM cells could be measured comparing co-cultures with NOXA-BCMA versus iNOXA-BCMA CAR T cells, no differences in the viability of other healthy BM cells were observed (**Fig. 4a,b**). This indicates there is no apparent toxicity to non-targeted BM cells. While NOXA was predominantly localized to granules in transduced T cells (**Fig. 2c** **and Extended Data** Fig. 3b) or NK cells (**Extended Data** Fig. 3c), it is possible that a portion of the introduced NOXA mis-localizes to -or leaks from-cytotoxic granules, resulting in toxicity to the T cells themselves. Therefore, we examined viability of transduced NOXA-BCMA versus iNOXA-BCMA CAR T cells during a 2-month in vitro culture period but did not observe differences in viability, while CD4:CD8 ratios and transcript expression remain comparable (**Fig. 4c**). Moreover, we examined presence of transduced NOXA-BCMA versus iNOXA-BCMA CAR T cells in blood and in the spleen of mice in a xenograft model that was previously outlined in **Fig. 3e-g**. This analysis reveals that addition of functional NOXA does not hamper in vivo persistence of BCMA CAR T cells (**Fig. 4d**).

**Figure 3.**
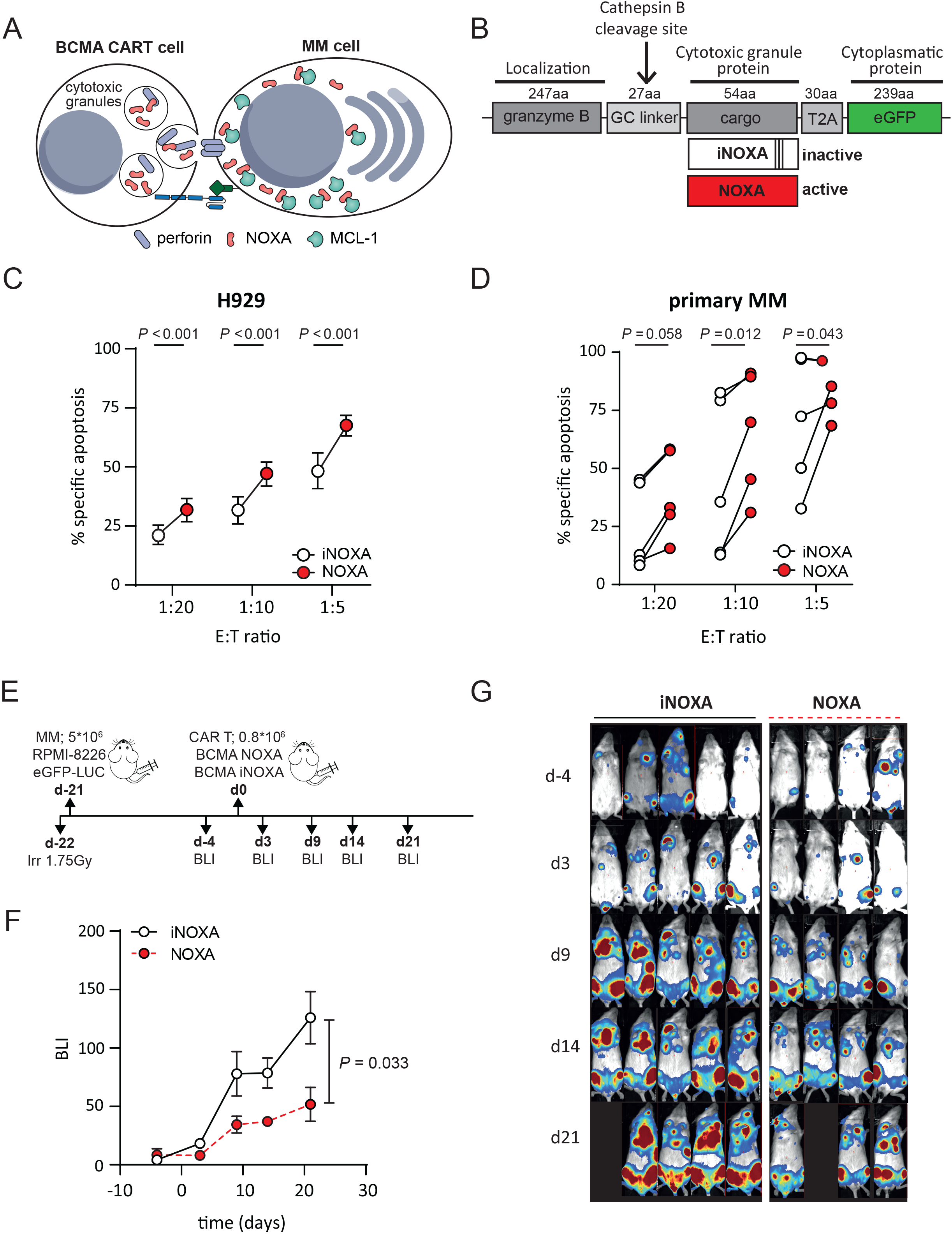
BCMA CAR T cells armed with NOXA show improved killing of MM cells. **A)** Graphical representation of optimized killing strategy by CAR T cells. **B**) Construct design with pro-apoptotic NOXA (active) or mutated NOXA (inactive) as cargo proteins fused to granzyme B. **C**) Specific apoptosis induced in H929 cells after 24 h of co-culture with NOXA-BCMA CAR T cells or iNOXA-BCMA CAR T cells at indicated E:T cell ratios. Values are average of 7 independent experiments with SEM. Statistical testing was performed using a two-way ANOVA followed by a Sidak’s multiple comparison test. **D**) Specific apoptosis induced in primary CD38^+^CD138^+^ MM cells after 48 hours of co-culture with NOXA-BCMA CAR T cells or iNOXA-BCMA CAR T cells at indicated E:T cell ratios. Dots represent different primary MM patient samples. Statistical testing was performed using a two-way ANOVA followed by a Sidak’s multiple comparison test. **E**) Experimental setup of xenograft mouse experiment where NSG mice are i.v. injected with RPMI8226-eGFP-Luc2 MM cells, followed by i.v. injection of indicated BCMA CAR T cells (0.8 x 10^6^, with a 1:1 CD4:CD8 ratio) 21 days later. **F**) Average bioluminescence intensity (BLI) (flux p/s) of mice treated with NOXA-BCMA or iNOXA-BCMA CAR T over time with SEM. Statistical testing was performed using a mixed effect model **G**) Corresponding BLI images of mice shown in (F).

**Figure 4.**
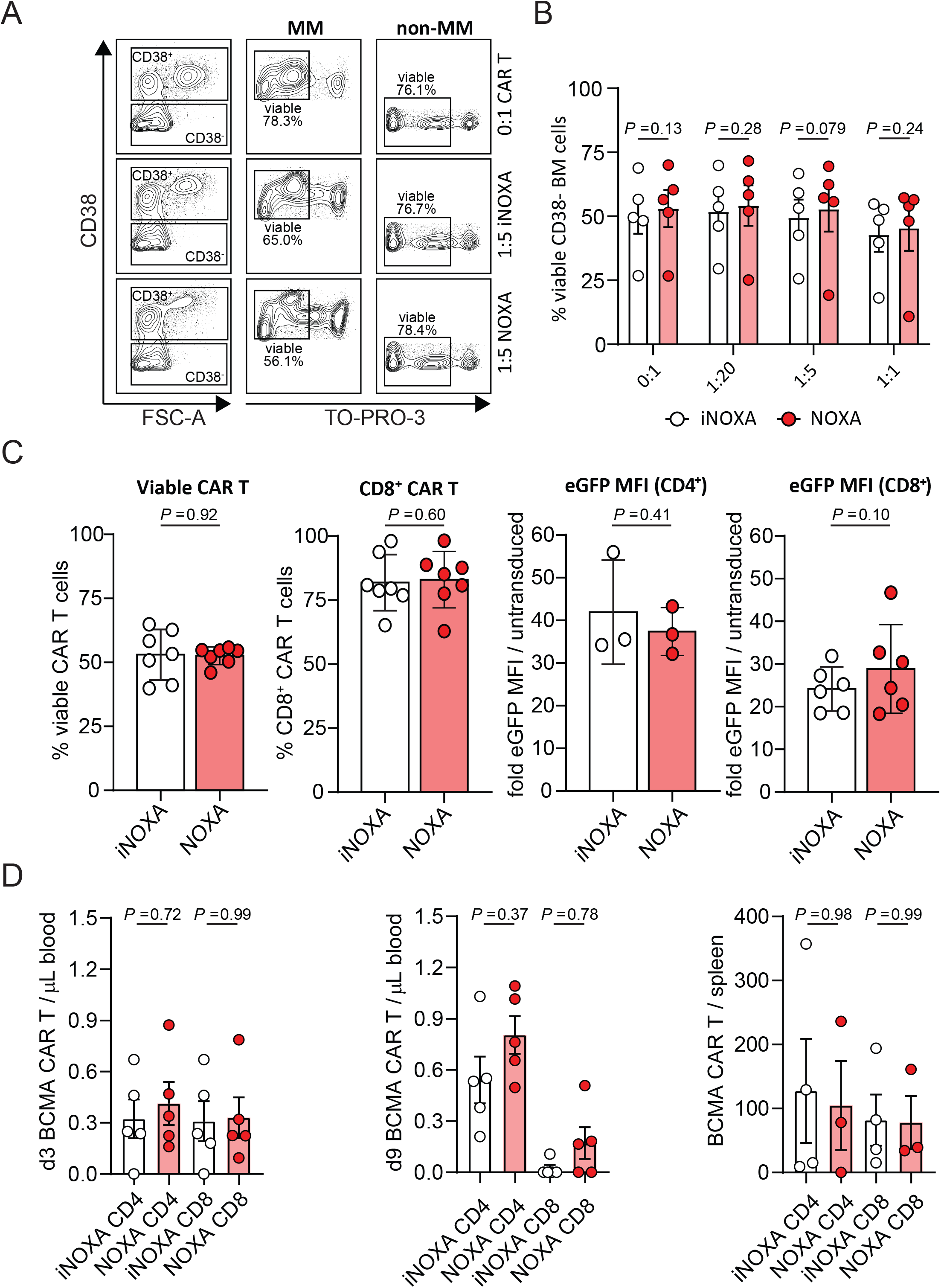
NOXA cargo does not impair viability of transduced BCMA CAR T cells and does not show off-target toxicity. **A)** Representative density plots showing cell viability (TO-PRO-3 staining) of primary bone marrow stromal cells (CD38^-^) or MM (CD38^+^) cells after 48 h of co-culture with iNOXA-BCMA or NOXA-BCMA CAR T cells in a 1:5 E:T cell ratio or without BCMA CAR T cells (0:1 E:T). **B**) Quantification of data shown in (A) with each dot representing a different MM patient sample co-cultured with iNOXA-BCMA or NOXA-BCMA CAR T cells at indicated E:T ratios for 48 h. Statistical testing was performed using a two-way ANOVA followed by a Sidak’s multiple comparison test. **C**) Plots showing total BCMA CAR T cell viability (TO-PRO-3-negative, left panel), percentage of CD8 (center left panel) and transcript (eGFP) expression in CD4^+^ (center right panel) and CD8^+^ (right panel) iNOXA-BCMA or NOXA-BCMA CAR T cells and cultured for 1 – 7 weeks after CAR T cell transduction. Every dot represents a separate experiment with SEM. Statistical testing was performed using one-way ANOVA. **D**) Number of CD4^+^ and CD8^+^ iNOXA-BCMA or NOXA-BCMA CAR T cells in blood at 3 (left panel) or 9 (center panel) days, or in the spleen (days 25-27, right panel), after i.v. injection in mice as outlined in figure 3E and F. Statistical testing was performed using one-way ANOVA, followed by multiple comparison testing.

## Discussion

Collectively, our data show that, by making use of the natural properties of granzyme B, we can direct pro-apoptotic proteins, such as NOXA, specifically into MM cells upon interaction with BCMA CAR T cells and subsequently promote apoptosis. As a consequence there is improved killing of cancer cells and diminished primary resistance. Importantly, resistance of tumor cells to undergo apoptosis after CAR T cell treatment is an immune escape mechanism that is not only confined to MM^2,3^. Therefore, tumor cell killing by CAR T cells in general should be improved to create an optimal CAR T cell therapy for more indications. By arming CAR T cells with NOXA we have demonstrated the possibility to improve killing of cancer cells dependent on MCL-1 expression for survival. However, this approach might not be sufficient for other tumor types that are MCL-1-independent. Pro-apoptotic proteins, besides NOXA, specifically inhibiting other pro-survival BCL-2 family proteins can be used in this setting. Alternatively, additional proteins that promote apoptotic or immunogenic cell death can be used to arm engineered T or NK cells and increase therapy options in a tailor-made fashion. Next to promoting cell death directly, other processes can be manipulated in targeted cancer cells by activating kinases or pathways, as shown previously using a synthetic enzyme-armed killer (SEAKER) CAR T cell strategy^29^. To examine CAR T cell behavior in more detail our strategy can also be employed using fluorescent molecules as cargo. This approach allows quantification of cargo transfer in relation to CAR T cell activation status, cellular interaction time and serial killing efficacy, and may provide clues for further optimization of CAR T cell technology.

## Methods

### Cell culture and chemicals

Cells were cultured in Dulbeccós Modified Eagle Medium (DMEM, Life Technologies) (Phoenix-Ampho, HEK293T), Iscovés Modified Dulbeccós Medium (IMDM, life Technologies) (OCI-Ly7), or RPMI 1640 GlutaMAX HEPES culture medium (Life Technologies) (L363, U266, MM1s, RPMI8226-GFP-Luc2, YT-Indy and pMM), supplemented with 10-20% fetal bovine serum (FBS, Sigma) and 100 µg/ml penicillin-streptomycin (p/s, Gibco/Life Technologies). For H929 cells 50 μM β-mercaptoethanol (Life Technologies) was added and for primary MM the medium was supplemented with 100 ng/ml human recombinant IL-6 (Tebu Bio) and 100 ng/ml human recombinant APRIL/TNFSF-13 (R&D systems). Human healthy donor peripheral blood mononuclear cells (PBMC) were isolated from buffy coats (Sanquin, Amsterdam, the Netherlands) using Ficoll-Paque according to the manufactureŕs protocol. PBMC were cultured in RPMI with 2.5% pooled AB+ human serum (IPLA-CSER, Innovative Research), 50 μM β-mercaptoethanol (Life Technologies) and 1% p/s. All primary MM samples were obtained after written informed consent, and protocols were approved by the local ethics committee of the University Medical Center, Utrecht.

### Apoptosis staining and flow cytometry

Assessment of cell viability was performed by staining with 20 nM TO-PRO-3 (Thermo Scientific) or with Fixable Viability Dye (FVD) eFluor506 or eFluor780 (eBioscience), followed by flow cytometric analysis (BD FACSCanto II or BD LSRFortessa, BD Biosciences). To determine the absolute amount of cells Flow count Fluorospheres were used (Beckman Coulter). In co-culture experiments, target cells were identified by flow cytometric surface staining with CD38-PE (Fisher) and CD138-PERCP-Cy5.5 (Biolegend) (DL-101) (primary MM cells), or by staining with CellTrace Violet (Invitrogen) (MM cell lines) prior to adding effector cells. BCMA CAR T cells were characterized by staining with CD4-Pacific Blue (Biolegend) (RPA-T4), CD8-PE/Cy7 (BD) (SK1), and biotinylated human BCMA (Bio-connect) with streptavidin-PE (Thermo Fisher). For intracellular staining, cells were fixed and permeabilized using BD Cytofix/Cytoperm (BD Biosciences), and stained with mouse anti-MCL-1 (Abcam) (Y37), rabbit anti-HA-TAG (CST) (C29F4), donkey anti-rabbit-IgG-Alexa Fluor 488 (Biolegend). Flow cytometry data analysis was performed using FlowJo.

### Generation of BCMA CAR T cells

The BCMA CAR construct (pBu-BCMA-CAR) was generated by cloning single chain variable fragments from anti-BCMA antibody into a pBullet vector containing a D8α-41BB-CD3-ζ signaling cassette. Phoenix-Ampho packaging cells were transfected with gag-pol (pHit60), env (P-COLT-GALV) and pBu-BCMA-CAR, using Fugene HD transfection reagent (Promega). Human PBMC were pre-activated with 30 ng/ml anti-CD3 (OKT3, Miltenyi) and 50 IU/ml IL-2 (Sigma) and subsequently transduced two times with viral supernatant in the presence of 6 ug/ml polybrene (Sigma) and 50 U/ml IL-2. Transduced T cells were expanded using 50 U/ml IL-2 and anti CD3/CD28 dynabeads (Thermo Fisher), and BCMA-CAR-expressing cells were selected by treatment with 80 µg/ml neomycin. T cells were further expanded using rapid expansion protocol as described elsewhere^30^. For transduction with our granzyme B – cargo – T2A - eGFP constructs, Gblocks (IDT) were cloned into lentiviral vector pCCL. Lentiviral particles were produced by transient transfection of the lentiviral vector and the packaging plasmids pRSV-Rev, pMDLg/pRRE, and pMD2-VSV-G to HEK293T cells using the CalPhos Mammalian Transfection Kit (Clontech Laboratories). Viral supernatants were filtered through 0.45 μm low-protein-binding filters, concentrated by ultracentrifugation at 20,000 × g for 2 h, resuspended in StemMACS (Miltenyi Biotec), and stored at −80°C. Previously transduced PBMC expressing a BCMA CAR were transduced with the granzyme B-cargo-expressing lentiviral vector and eGFP-positive BCMA CAR T cells were subsequently sorted (Sony MA900) and expanded on rapid expansion protocol.

### Animal model

NOD.Cg-PrkdcscidIl2rgtm1Wjl mice were ordered (Charles River) and temporarily housed in the Central Animal Facility of Utrecht University during the experiments. Experiments were conducted per institutional guidelines after obtaining permission from the local ethical committee, and performed in accordance with the current Dutch laws on animal experimentation. Mice were housed in individually ventilated cage (IVC) system to maintain sterile conditions and fed with sterile food and water. After irradiation, mice were given the antibiotic ciproxin in the sterile water throughout the duration of the experiment. Female mice were randomized with equal distribution among the different groups, based on tumor size (measured with Bioluminescence Imaging (BLI) on day −1) into 5 mice/group. Age and weight of mice was comparable between groups. Adult NSG mice (6-9 weeks old) received sublethal total body irradiation (1,75 Gy) on day −22 followed by intravenous injection of 5*10^6^ RPMI-8226-luciferase tumor cells on day -21, and received 1 intravenous injection of 0.8*10^6^ NOXA BCMA CAR T cells or iNOXA BCMA CAR T cells on day 0. Together with the CAR T cell injection, all mice received 0.6 × 10^6^ IU of IL-2 (Proleukin; Clinigen) in 100 µl incomplete Freund’s adjuvant (IFA). Mice were monitored at least twice a week for any symptoms of disease (sign of paralysis, weakness, and reduced motility), weight loss, and clinical appearance scoring (scoring parameter included hunched appearance, activity, fur texture, and piloerection). The humane endpoint was reached when mice showed the aforementioned symptoms of disease or experienced a 20% weight loss from the initial weight (measured on day 1). During the experiment tumor growth was monitored weekly by BLI measurement after intraperitoneal (IP) Luciferin (Promega) injection.

### Exogenous delivery of NOXA

In order to determine the effect of NOXA mediated killing, the pore forming Streptolysin O (S5265-25KU, Sigma) was used to facilitate entry of exogenous NOXA into target cells. SLO was activated with 10 mM DTT for 20 minutes at RT and subsequently diluted in serum-free RPMI. Target cells were incubated with SLO and synthetic NOXA or NOXA-TAMRA for 30 minutes at 37 °C, after which FBS-containing medium was added to inactivate the SLO. After 24h, apoptosis staining was performed to measure target cell viability using flow cytometry.

### Immunoblotting

For western blot analysis, cells were lysed in buffer containing 1% NP-40 and proteins were separated using SDS-PAGE (Mini-PROTEAN® TGX™ Precast Gels, Bio-Rad), transferred to low fluorescence PVDF membranes (Bio-Rad), blocked in phosphate-buffered saline (PBS) containing 2% non-fat dry milk, and stained using the following antibodies: mouse anti-α-tubulin (Cell signaling technology) (DM1A), mouse anti-NOXA (Abcam) (114C307.1), rabbit anti-MCL-1 (Abcam) (Y37), goat anti-mouse-680RD, and goat anti-rabbit-800CW (LI-COR Biosciences). To enrich for MCL-1 binding protein immunoprecipitation was preformed using the Dynabeads Protein G IP Kit (Invitrogen) following the manufacturer’s protocol. Infrared imaging was used for detection (Odyssey Sa; LI-COR Biosciences). Analysis and quantification were performed using LI-COR Image Studio and ImageJ 1.47V software.

### Microscopy

To visualize mScarlet, mNeongreen or NOXA, CAR-T cells were placed on coverslips containing 0.1% poly-l-lysine. Consequently, cells were fixed using 4% paraformaldehyde. For intracellular staining cells were blocked for 1 hour using a blocking buffer consisting of 2% BSA and 0.1% saponin in PBS, followed by 1-hour incubations with anti-LAMP-1 (BD or CST) (H4A3 or D2D11) or anti-HA tag (CST) (C29F4), in blocking buffer. After washing, the coverslips were mounted using ProLongTM Gold with DAPI (Invitrogen). For visualizing transfer of mScarlet or mNeongreen from BCMA CAR T cells into MM cells, target cells were stained using VybrantTM DiO Dye prior to incubation with the effector cells. Cells were imaged using a 63 x oil lens (630 x total magnification) on a confocal microscope (Zeiss LSM710 (fixed) or Stellaris 5, Leica Microsystems (live imaging)). ImageJ was used to analyze the images.

### Statistical analysis

Statistical analysis was performed using GraphPad Prism version 8.3. Unpaired groups were compared with a Student’s t-test. For comparison of more than two groups, one-way or two-way ANOVA tests were used.

## Acknowledgments

The authors thank the support facilities of the University Medical Center Utrecht. Synthetic NOXA and NOXA-TAMRA were kindly provided by Prof. dr. H. Ovaa, Department Cell and Chemical Biology, Leiden University Medical Centre, Leiden, The Netherlands. This work was financially supported by research grants from the Dutch Cancer Foundation (KWF)/Alpe d’HuZes Foundation (11270 and 13058 to V.P, and 14459 to M.C.) The funding agency played no role in the design, reviewing, or writing of the manuscript.

---

**Extended Data Figure 1.**
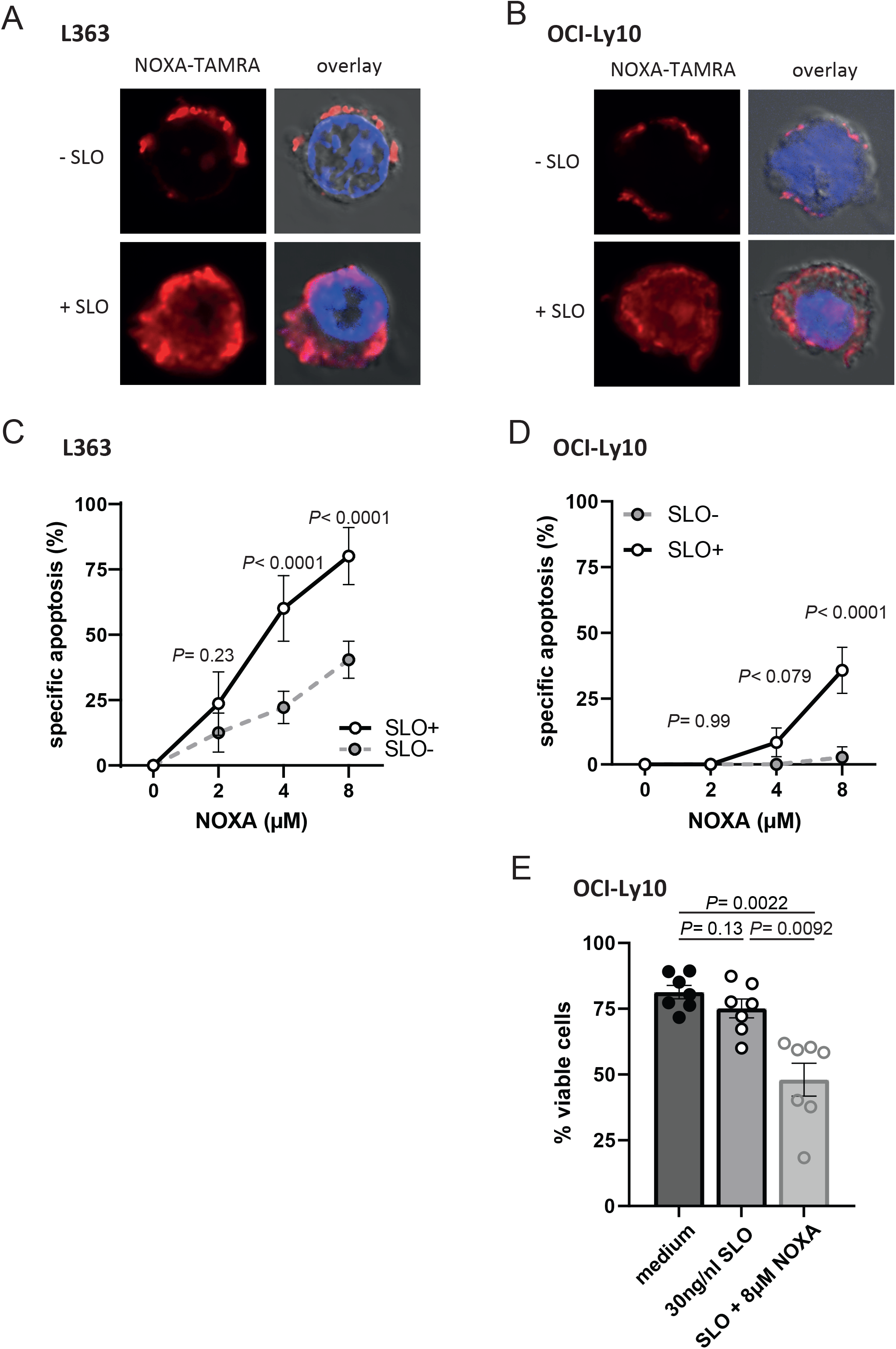
Exogenous NOXA induces apoptosis in L363 and OCI-Ly10 cells. **A,B**) Representative confocal image (630 x oil magnification) showing A) L363 MM cells incubated with 10 µM synthetic NOXA-TAMRA with or without 15 ng/ml streptolysin O (SLO) and B) OCI-Ly10 DLBCL cells with 10 µM synthetic NOXA-TAMRA with or without 30 ng/ml streptolysin O (SLO). **C**) Apoptosis induced by synthetic NOXA in L363 MM cells treated with 15 ng/ml SLO or without SLO, analyzed by flow cytometry with viability dyes TO-PRO-3 and DiOC6(3). Shown are averages of 5 biological replicates with SEM. Statistical analysis was performed using a two-way ANOVA followed by multiple comparison testing. **D**) Apoptosis induced by synthetic NOXA in OCI-Ly10 DLBCL cells treated with 30 ng/ml SLO or without SLO, analyzed by flow cytometry with viability dyes TO-PRO-3 and DiOC6(3). Shown are averages of 5 biological replicates with SEM. Statistical analysis was performed using a two-way ANOVA followed by multiple comparison testing. **E**) Percentage of viable (DiOC6(3)^+^TO-PRO-3^-^) OCI-Ly10 DLBCL cells treated with 30 ng/ml SLO, with or without 8 µM synthetic NOXA and analyzed by flow cytometry. Statistical analysis was performed using a one-way ANOVA with Geisser-Greenhouse correction.

**Extended Data Figure 2.**
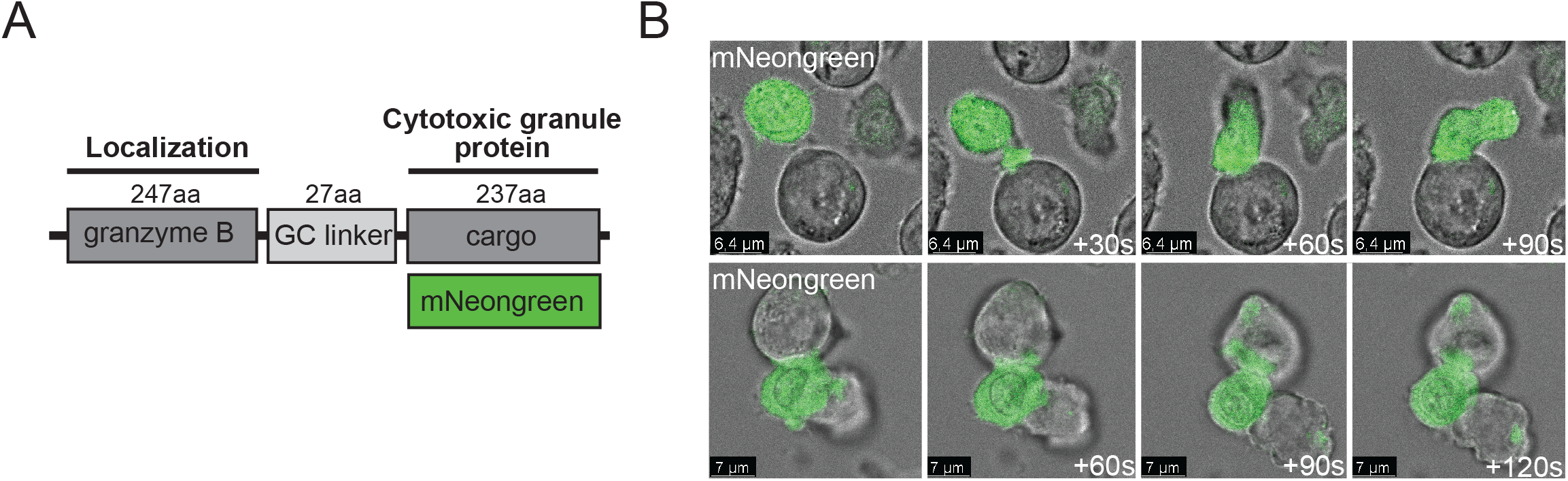
Visualizing fluorescent cargo from BCMA CAR T cells to targeted MM cells. **A)** Construct design with fluorescent mNeongreen fused to granzyme B. **B**) Brightfield images combined with fluorescent intensity of mNeongreen (green) of H929 MM cells pre-treated with 10 µM Q-VD-OPh and co-cultured with BCMA CAR T cells transduced with construct shown in (G) in a 1:1 E:T ratio (confocal microscopy 630 x oil magnification). Images were taken with a 30 second interval.

**Extended Data Figure 3.**
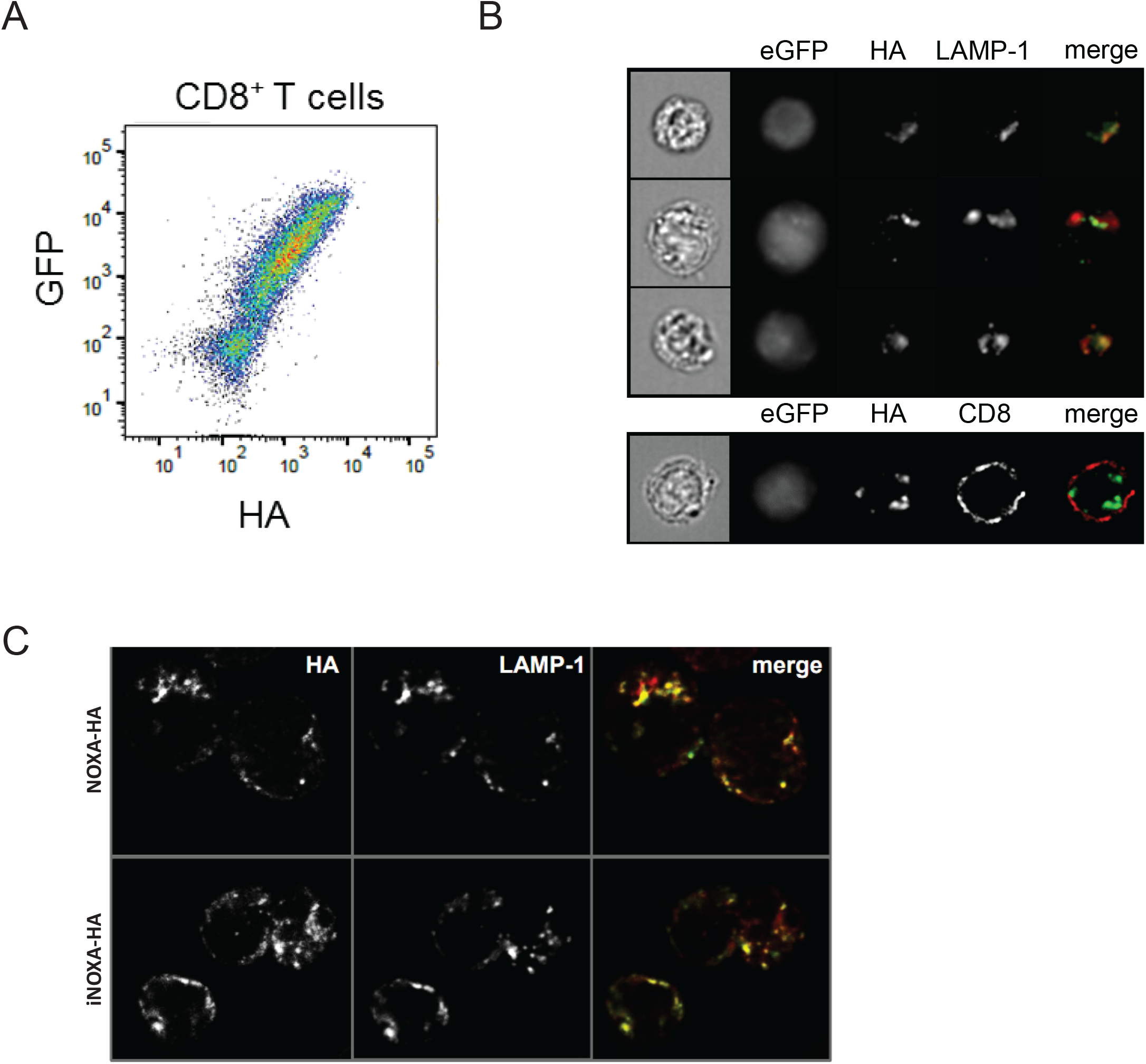
NOXA localizes to LAMP-1-postive cytotoxic granules in transduced primary CD8^+^ cells and NK cells. **A)** Representative flow cytometry staining of CD8^+^ BCMA CAR T cells lentivirally transduced with a GzB-NOXA-T2A-eGFP construct (NOXA-BCMA CAR T cells) as shown in Figure 3B and stained with an anti-HA antibody. **B**) Representative image-based flow cytometry image captures by ImageStream at 60x magnification of transduced CD8^+^ BCMA CAR T as in (A) and stained with antibodies against HA and LAMP-1 or CD8. **C**) Representative immuno-fluorescent staining of YT-Indy cells NK cells lentivirally transduced with GzB-NOXA-T2A-eGFP or GzB-iNOXA-T2A-eGFP constructs as shown in Figure 3B for late endosomal marker LAMP-1 and HA (to visualize the HA-tag coupled to NOXA or iNOXA), and analyzed by confocal microscopy (630 x oil-magnification).

**Extended Data Figure 4.**
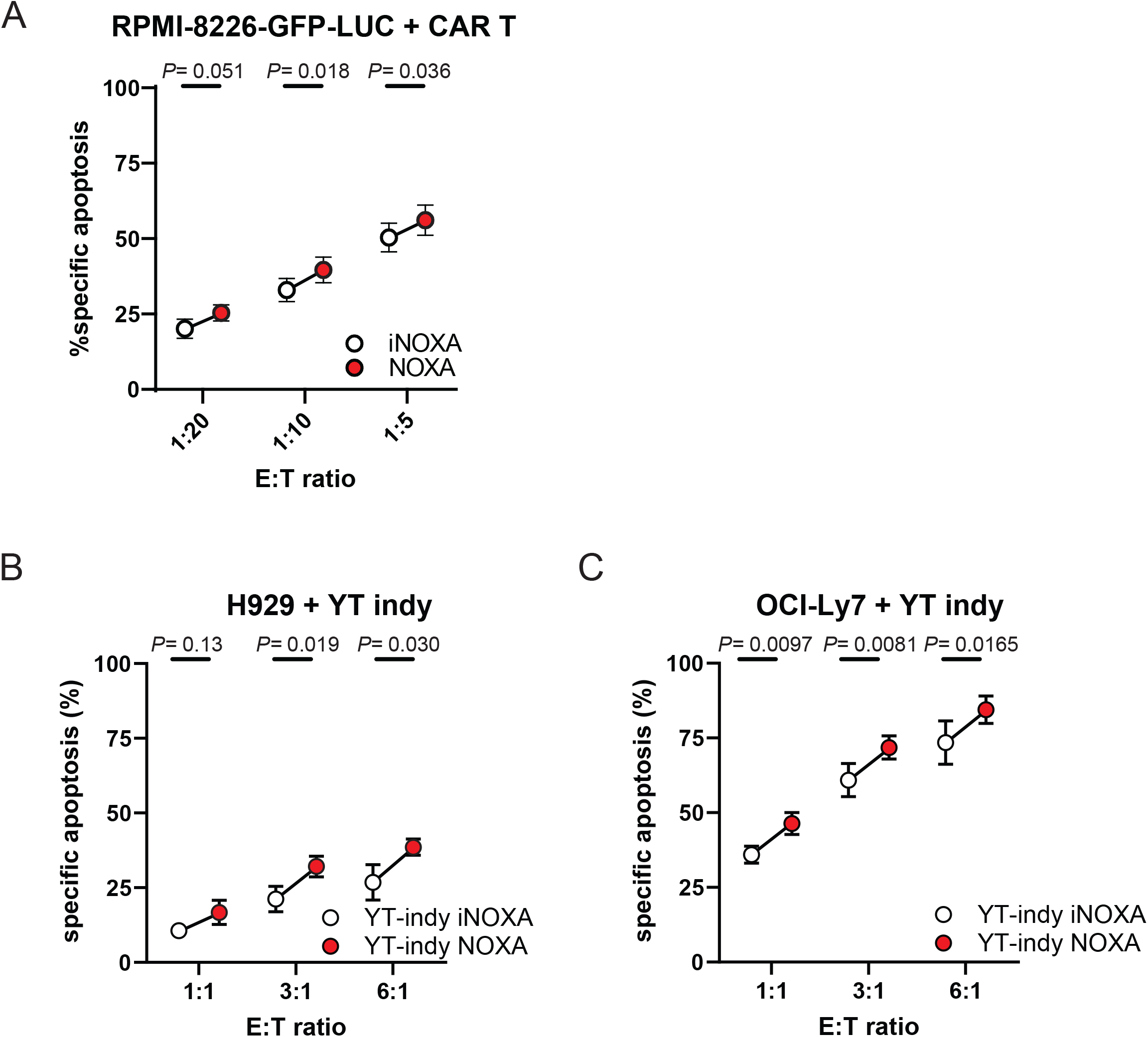
BCMA CAR T cells or NK cells armed with NOXA show improved killing of MM or DLBCL cells. **A)** Specific apoptosis as measured by TO-PRO-3 expression using flow cytometry induced in RPMI-8226-GFP-Luc2 cells after 24 h of co-culture with iNOXA-BCMA or NOXA-BCMA CAR T cells at indicated E:T cell ratios. Values are average of 4 independent experiments with SEM. Statistical testing was performed using a two-way ANOVA followed by a Sidak’s multiple comparison test. **B**) Specific apoptosis induced in H929 MM cells after 4 h of co-culture with YT-Indy NK cells lentivirally transduced with GzB-NOXA-T2A-eGFP or GzB-iNOXA-T2A-eGFP constructs as shown in Figure 3B at indicated E:T cell ratios. Values are average of 3 independent experiments with SEM. Statistical testing was performed using a two-way ANOVA followed by a Sidak’s multiple comparison test. **C**) Specific apoptosis induced in OCI-Ly7 DLBCL cells after 4 h of co-culture YT-Indy cells NK cells lentivirally transduced with GzB-NOXA-T2A-eGFP or GzB-iNOXA-T2A-eGFP constructs as shown in Figure 3B at indicated E:T cell ratios. Values are average of 3 independent experiments with SEM. Statistical testing was performed using a two-way ANOVA followed by a Sidak’s multiple comparison test.

